# In Situ Transcriptome Accessibility Sequencing (INSTA-seq)

**DOI:** 10.1101/722819

**Authors:** Daniel Fürth, Victor Hatini, Je H. Lee

## Abstract

Subcellular RNA localization regulates spatially polarized cellular processes, but unbiased investigation of its control *in vivo* remains challenging. Current hybridization-based methods cannot differentiate small regulatory variants, while *in situ* sequencing is limited by short reads. We solved these problems using a bidirectional sequencing chemistry to efficiently image transcript-specific barcode *in situ*, which are then extracted and assembled into longer reads using NGS. In the *Drosophila* retina, genes regulating eye development and cytoskeletal organization were enriched compared to methods using extracted RNA. We therefore named our method In Situ Transcriptome Accessibility sequencing (INSTA-seq). Sequencing reads terminated near 3’ UTR *cis*-motifs (e.g. *Zip48C, stau*), revealing RNA-protein interactions. Additionally, *Act5C* polyadenylation isoforms retaining zipcode motifs were selectively localized to the optical stalk, consistent with their biology. Our platform provides a powerful way to visualize any RNA variants or protein interactions *in situ* to study their regulation in animal development.

---

Differential hybridization and cloning of spatially separated mRNAs obtained from distinct dissected regions of *Xenopus* oocytes were reported thirty years ago (1–4). It revealed the role of asymmetrically localized mRNAs and 3’ Un-Translated Regions (UTRs) in development, and similar observations were made in other systems (5). Despite advances in molecular methods (6) unbiased investigation of mRNA localization, 3’ UTR isoforms, and RNA-binding proteins (RBPs) *in situ* remains challenging.

In *Drosophila* embryos a majority of mRNAs examined by fluorescent *in situ* hybridization (FISH) have been shown to exhibit distinct subcellular localization patterns (7). During development of the *Drosophila* retina, epithelial cells organize into a precisely ordered structure composed of approximately 800 hexagonal facets known as ommatidia. Each ommatidium consists of several distinct cell types including a core of eight photoreceptor cells capped by four cone cells and surrounded by two large primary (1°) pigment cells, along with 2° and 3° pigment cells and mechanosensory bristle cells in the perimeter (**Fig. 1a**) (8–10). The emergence of the precise cellular arrangement in ommatidia has been used to investigate basic principles involved in pattern formation and morphogenesis (11), making it an ideal system for studying RNA localization associated with cellular geometry and tissue patterning.

**Fig. 1.**
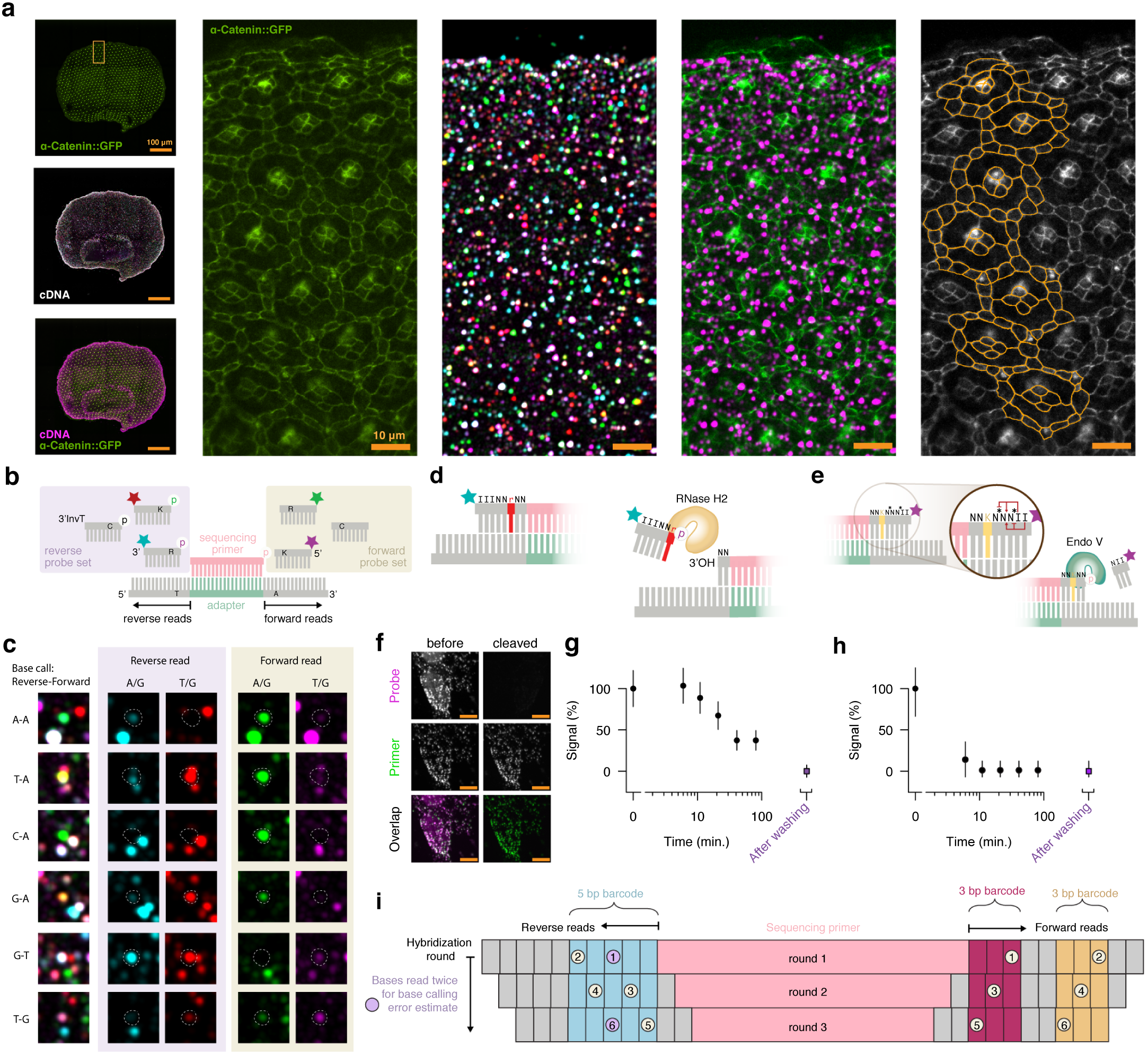
Paired-end Ribonucleotide-Inosine Cleaved *K*-mer Ligation (PRICKLi) chemistry. **a)** Top first column: a third instar *Drosophila* retina with GFP-tagged *α*-catenin. Middle first column: *In situ* cDNA sequencing cycle one. Bottom first column: Fluorescent probe hybridization to cDNA amplicons (magenta). Second column: a close-up view of the *α*-catenin::GFP signal. Third column: PRICKLi sequencing. Fourth column: fluorescent sequencing primer hybridization (magenta). Last column: cell segmentation (orange) using GFP-tagged *α*-catenin signal (gray). **(b)** PRICKLi uses two colors to discriminate 4 bases at each end simultaneously, interrogating the third base from the ligation junction using mixed bases (K or R). **(e)** Examples of two-color reverse and forward base calling (dotted circles). The two fluorophores at each end represents either A/G or T/G, enabling one to determine the base identity. When a fluorescent spot disappears during sequencing, the amplicon base is called as C. **(d)** Reverse probes incorporate K or R ribonucleotides for RNase H2 cleavage, generating a free 3’ OH end. **(e)** Forward probes incorporate two inosines (I) at the 5’ end along with phosphorothioate modifications (*) for site-specific Endonuclease V cleavage. **(f)** cDNA amplicons in cultured cells were hybridized to sequencing primers (green), followed by ligation of PRICKLi probes (magenta). After one hour, all cDNA amplicons with sequencing primers were labeled by PRICKLi fluorophores. The loss of PRICKLi fluorescence relative to the sequencing primer fluorescence was used to determine the kinetics of **(g)** RNase H2 and **(h)** Endonuclease V cleavage. **(i)** A schematic of six PRICKLi imaging cycles. In each round, a sequencing primer is hybridized, followed by two cycles of ligation and imaging with a single cleavage step in-between. Using recessed sequencing primers for round 2 and 3, six contiguous bases are interrogated in the reverse direction, while six bases with a two base gap are determined in the forward direction.

Multiple methods can measure the transcript abundance *in situ* (12–22). Yet, it remains difficult to spatially map regulatory elements and RBPs in a systematic and unbiased manner. One approach to mapping regulatory motifs, nucleotide variants, or mRNA isoforms is to synthesize and sequence cDNA directly in the tissue (12). Here we hypothesized that if cDNA termination events were to be localized *in situ* it could even be possible to map nucleotide signatures associated with mRNA-RBP cross-linking (6) within intact cells. To map both cDNA synthesis start and stop sites *in situ*, however, a mated-pair sequencing approach with a sufficiently long read length is necessary.

Here we report a novel bidirectional sequencing chemistry called PRICKLi (Paired-end Ribonucleotide-Inosine Cleaved *K*-mer Ligation) that sequences two bases simultaneously from both ends of the cDNA fragment. It allows for paired-end *in situ* reads in one imaging cycle and reduces the number of 12-base barcode imaging cycles to six. To generate full-length cDNA reads, we decoupled imaging from sequencing by reading transcript-associated barcodes *in situ*, followed by cDNA amplicon extraction and Next-Generation Sequencing (NGS). The shared barcode sequences are then used to map assembled NGS reads to a reference tissue atlas bearing barcode coordinates (14, 22).

Together, these advances enable spatial mapping of major regulatory sequences, factors, and RNA-protein interactions involved in subcellular mRNA localization, especially in 3’ UTRs with alternative cleavage or polyadenylation (APA) isoforms. We refer to this method as *in situ* transcriptome accessibility sequencing (INSTA-seq).

## Results

### Paired-end *in situ* sequencing and decoupling of read length from imaging

Unlike sequencing-by-synthesis (SBS) that extends the sequencing primer in one direction (5’ to 3’), PRICKLi interrogates both ends of the sequencing primer simultaneously using sequencing-by-ligation (SBL) (**Fig. 1b**). Fluorescently labeled probes containing mixed bases (K: G/T, R: A/G) along with a dark probe (C) utilize the co-localization pattern of two colors from each amplicon to make a base call (**Fig. 1c**). To cleave the terminal fluorophore, reverse PRICKLi probe incorporates ribonu cleotides (rK, rR, or rC) that are nicked by RNase H2 (**Fig.** 1d) at the 5’ side (23), while the forward PRICKLi probe incorporates inosine for Endonuclease V-dependent cleavage (24). Because the cleavage position of Endonuclease V is variable, phosphorothioate modifications are used to block undesired cleavage sites (**Fig. 1e**) (25).

After testing the efficiency of ligation and cleavage on cultured cells and beads (**Fig. S1**), we sequenced cDNA amplicons in the *Drosophila* retina (**Fig. 1c**), demonstrating complete cleavage of the terminal fluorophores by RNase H2 or Endonuclease V after 40 or 10 minutes, respectively, prior to the next ligation cycle (**Fig. 1f-h**). By cycling through ligation, imaging, and cleavage, we sequenced twelve bases using four fluorescent oligonucleotides in six imaging cycles (**Fig. 1i**).

We then performed sequential ligation-cleavage cycles across multiple *Drosophila* retinas to obtain transcript-associated optical barcodes. Dissected retina specimens were mounted whole (3-6 per well; 9 wells) onto a cover glass (**Fig. 2a**). After fixation, genomic DNA digestion (**Fig. S2**), RT, circularization, and rolling circle amplification (RCA), we sequenced 12 bases from cDNA amplicons on a confocal microscope (40 × magnification, 9-12 tiles per retina, 90-200 *z*-stacks with a 500-nm step size) (**Fig. 2a**). Because PRICKLi uses dual color base calling, individual amplicons were segmented and registered in 3D across four channels to account for optical aberration, displacement, or variable ligation. Subsequently, base calls from each amplicon were assigned by clustering (**Fig. 2b-c**). For segmentation we developed a set of wavelet filters (**Fig. 2d**) that extract the energy from fluorescent spots with a defined period (e.g. single amplicon size) to suppress background noise (**Fig. S2**). Because cytosine-cytosine base calls are inferred by the disappearance and reappearance of fluorescence across imaging cycles, such calls are only made after 3D registration (**Fig. 2d**). Non-rigid 3D point set registration by coherent point drift enables (**Fig.** 2e) tracking of amplicons across cycles despite optical aberrations or sample displacement. The end result is a full 3D reconstruction of transcript-associated barcode reads in the retina whole-mount (**Fig. 2f**).

**Fig. 2.**
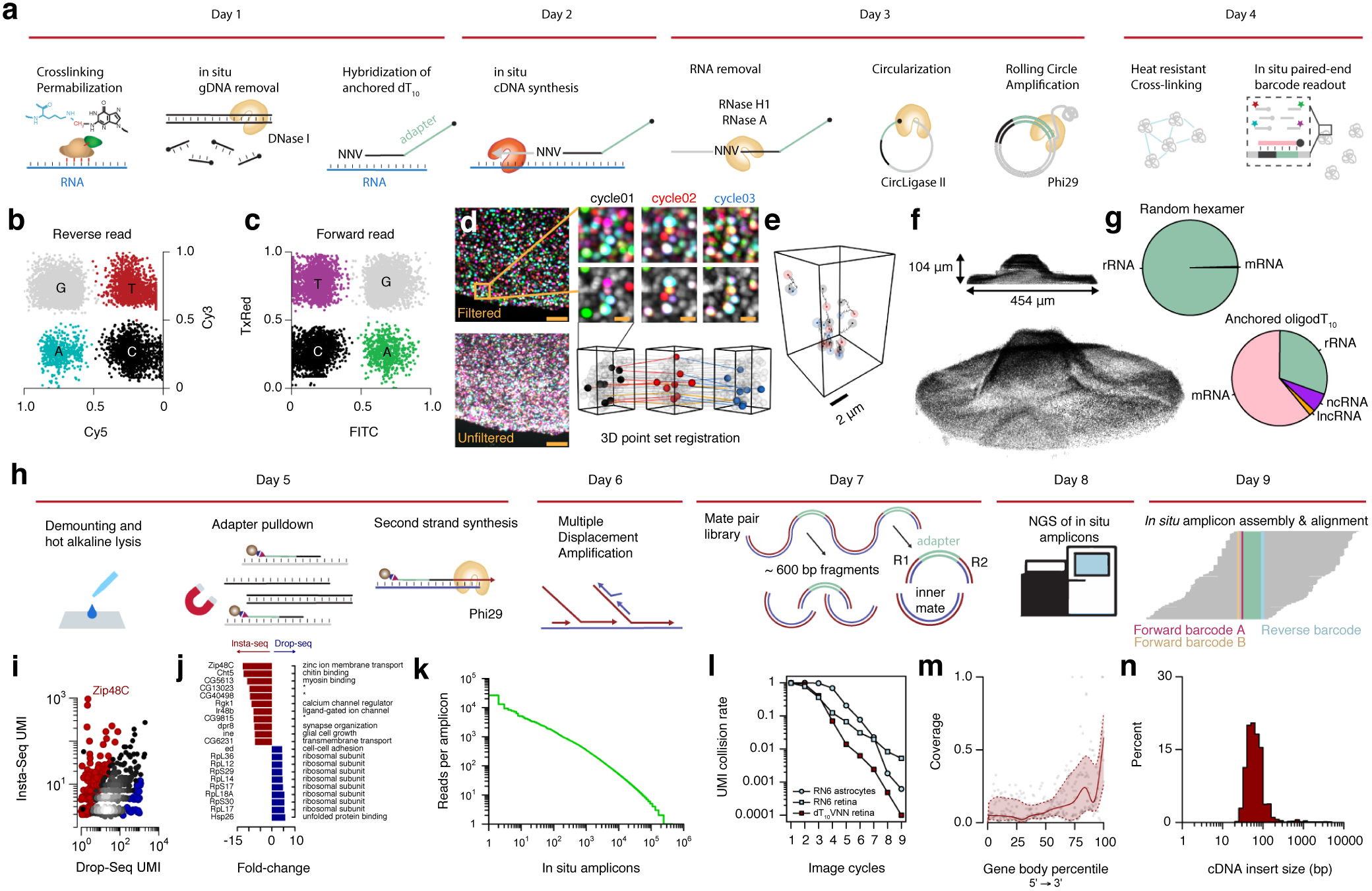
Compressing transcriptome-wide interrogation into six imaging cycles decouples read length from imaging time. **a)** Workflow of *in situ* sequencing cDNA library preparation. Clustering of detected amplicons at each cycle into b) reverse and **c)** forward reads. **d)** To reduce base calling errors due to high amplicon density or optical aberration, cDNA amplicons were segmented and registered in 3D to generate corresponding indexes. A close up on a 10 × 10 x 14-µm volume comprised of 101 amplicons is shown, along with eight amplicons highlighted between cycles. Orange lines denotes cDNA amplicons that disappears on cycle 2 and reappears in cycle 3 (‘C’ base call). Scale bars: 10 µm and 1 µm. **e)** Estimated non-rigid displacement of the highlighted eight amplicons from cycle 1 (black), cycle 2 (red) and cycle 3 (blue). **f)** Complete 3D reconstruction of detected *in situ* amplicons in *Drosophila* retina. **g)** The yield of different RNA species using either random hexamer or anchored oligo(dT)10 in the third instar *Drosophila* retina. **h)** Workflow of NGS library preparation for sequencing *in situ* amplicons with Illumina paired-end sequencing. The retina sample is demounted and lysed followed by hybridization of a biotinylated sequencing primer and pull-down using streptavidin magnetic beads. The second strand is synthesized, followed by MDA and Tn5 tagmentation. Multiple tissue samples are indexed using i5/i7 indexes and P5/P7 adapters for subsequent NGS. **i)** The correlation of average UMI count for individual genes between INSTA-seq and DROP-seq (INSTA-seq enriched genes in red; DROP-seq enriched genes in blue). **j)** Top ten differentially enriched genes in INSTA-seq (red) vs. DROP-seq (blue) shows preferential depletion of ribosomal protein genes in INSTA-seq. **k)** Histogram of NGS reads per amplicon. **l)** The UMI collision rate as a function of number of imaging cycles, RT primers (dark red: oligodT10; light blue: random hexamer), and tissue types (squares: *Drosophila* retina; circles: human immortalized astrocytes). **m)** Gene body coverage of INSTA-seq across replicates. **n)** Histogram of the cDNA insert size in the *Drosophila* retina.

Previously, we observed 10-40% of cDNA reads were generated from mRNA when using random hexamer for RT (12). In contrast, we observed over 99.6% rRNA in retina from third instar larva (**Fig. 2g**). By using a short NNV-oligo(dT)10 primer instead, we enriched mRNA reads by 155-fold (61%), in addition to long (2%) and short non-coding RNAs (6.5%) (**Fig. 2g**). In total we detected 820 unique protein-coding genes.

Circular cDNAs generate paired sequences corresponding to reverse transcription (RT) termination and start sites. To characterize internal errors caused by cross-linked nucleotides (26), we used NGS to generate longer cDNA reads. Briefly, PRICKLi-sequenced tissues were lysed, followed by hybridization-based enrichment of cDNA amplicons. After second strand synthesis, multiple displacement amplification (MDA), tagmentation, and tissue multiplexing, we generated mated-pair reads on MiSeq (**Fig. 2h**). Compared to DROP-seq of the *Drosophila* eye disc (**Fig. 2i**) (27), INSTA-seq enriched for genes involved in neurogenesis, axon development, and cytoskeleton organization (**Table S1-2**). Interestingly, INSTA-seq detected relatively few ribosomal subunit transcripts, although they comprised the largest fraction in the DROP-seq data (**Fig. 2j**). The reason for this is not clear but suggests that certain biologically relevant transcripts are more accessible by our method. The number of NGS reads per amplicon (443,304 UMIs in total) followed a geometric distribution with no apparent saturation (**Fig. 2k**), suggesting that deeper sequencing could recapture more *in situ* amplicons. Since the number of amplicons determines the frequency of non-unique barcodes (‘collision rate’), both tissue size and amplicon density are important considerations. Although each additional PRICKLi imaging rounds can decrease the collision rate by 28% per imaging round (95% CI: 13-63%) (**Fig. 2l**), the utility of longer UMIs needs to be weighed against increased imaging time. In the end, we settled on a collision rate of 0.62%, which is compatible with six rounds of 3D imaging across *Drosophila* whole-mount retinas in a single day.

### INSTA-seq for unbiased profiling of *cis*-acting RNA localization elements

To identify putative 3’ UTR *cis* motifs, we asked whether synthesized cDNA synthesis terminated pre-maturely adjacent to known *cis* motifs. Indeed, we found that transcript body coverage was skewed towards the end of 3’ UTRs, suggesting incomplete RT (**Fig. 2m**) with the cDNA size ranging from 8-nt to 4750-nt; however, most amplicons were <100-nt (mean: 86-nt, s.d.: 229-nt) (**Fig. 2n**). In fact, only 1.74% of cDNAs reached the coding region, while 15% terminated abruptly in the 3’ UTR with a terminal single-nucleotide mismatch (**Fig. 3a**). Single-nucleotide mismatches were more frequent at the 3’ end compared to the 5’ end of the cDNA (*χ*^2^ 274.96, *p*-value < 10^−4^) (**Fig. 3b**), and internal mismatches were rare (< 0.06%) with the majority being deletions rather than substitutions. Most mismatches near the 3’ end were due to a single nucleotide (66.22%), and no more than eight nucleotides long (1.73%) (**Fig. 3c**). Unlike cytosine-tailing by M-MuLV reverse transcriptase (28), the incorporation of cytosine or homopolymer tailing was infrequent, while consecutive mismatches reflected the AU-rich elements in the 3’ UTR (**Fig. 3c**). These results suggest that short cDNA fragment are truncated cDNAs rather than cDNAs synthesized from fragmented RNAs.

**Fig. 3.**
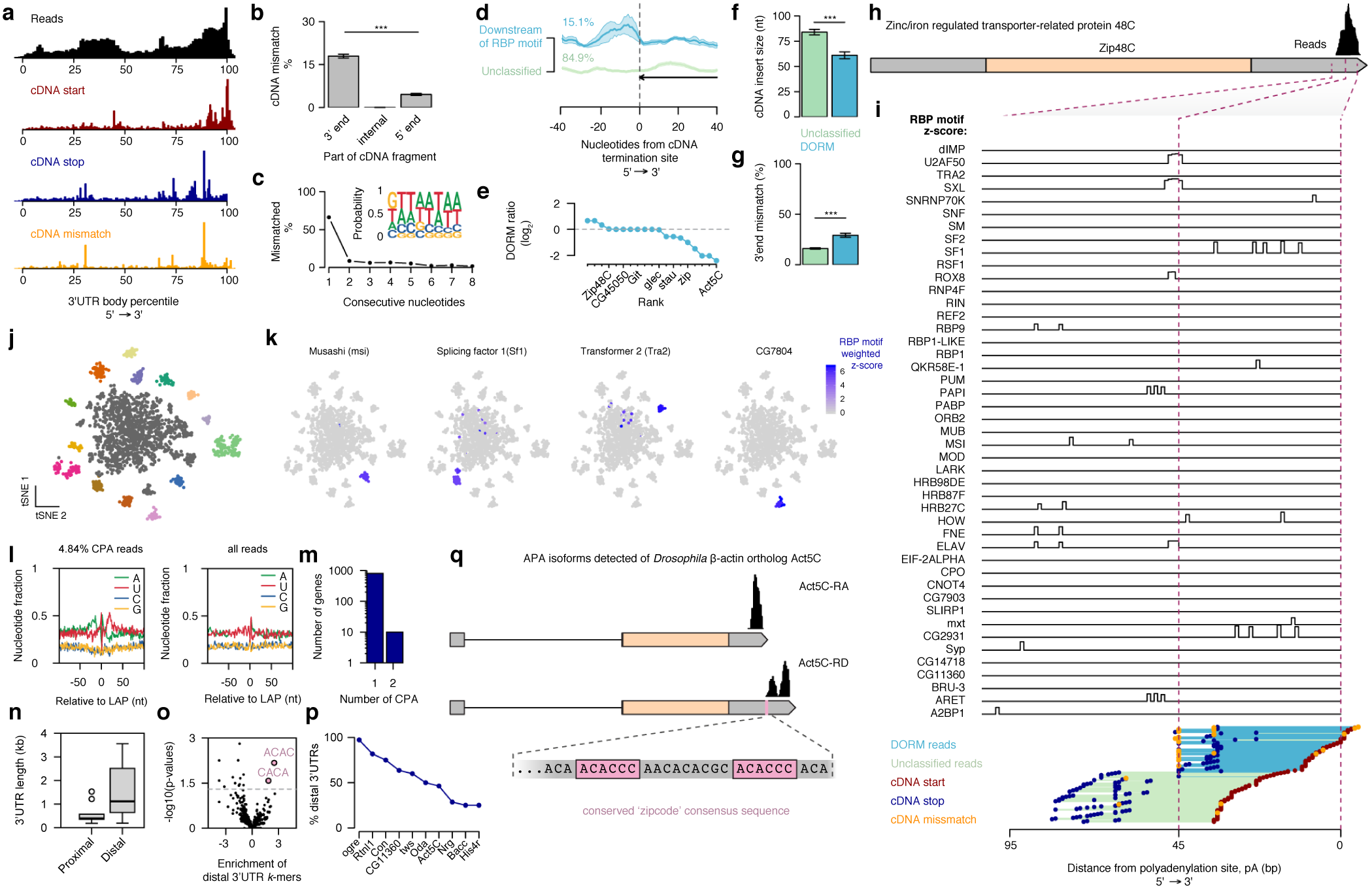
INSTA-seq identifies 3’ UTR *cis*-acting elements and putative *trans*-acting partners. **a)** Top black: Coverage across the 3’ UTR body percentile. Dark red: Histogram of cDNA synthesis starting sites. Dark blue: Histogram of cDNA termination sites. Orange: Histogram of cDNA mismatch sites. **b)** Percent of cDNA mismatches as a function of position within the cDNA. **c)** The number of consecutive mismatched nucleotides in cDNAs with a terminal mismatch. Inset: Probability of mismatched nucleotides from the cDNA 3’ end. **d)** Average bedtracks of the RBP motif probability near the cDNA termination site. cDNAs were clustered into reads exhibiting an increase in the RBP motif probability adjacent to the cDNA termination site (blue: Downstream-Of-RBP-Motif or DORM) or unclassified (green). **e)** Log-ratio of DORM reads vs. reads that pass through the DORM site (non-DORM reads) for nineteen genes in the cluster. **f)** DORM clustered reads show a shorter cDNA insert size compared to non-DORM reads. **g)** DORM reads show a higher percentage of 3’ end mismatches compared to non-DORM reads. **h)** Structure and read coverage of *Zip48C* gene. **i)** A close-up view of individual bedtracks of RBP motif match z-scores in the *Zip48C* 3’ UTR (bottom: DORM cluster reads in blue and non-DORM cluster reads in green). **j)** Dimensionality reduction and clustering of DORM reads. **k)** Weighted *z*-score of individual RBP motifs show cluster-specific RBP motifs. **l)** Nucleotide composition around the last aligned position LAP (set to position 0) for Cleavage and Polyadenylation site-supporting (CPA) reads (left) and all reads (right). CPA reads exhibit U-rich CstF-binding sites 20-bp downstream of the LAP. **m)** Number of genes with one or two CPAs. **n)** Boxplot of proximal vs. distal 3’ UTR length in kilobase pairs. **o)** Volcano plot with log2 fold-change for tetramers in the distal 3’ UTR compared to the proximal 3’ UTR across 10 genes with proximal and distal isoforms. Highlighted in pink are CA-rich tetramers enriched in distal but not proximal 3’ UTR isoforms for *Act5C*. **p)** Genes ranked according to the percentage of transcripts with the distal 3’ UTR. **q)** Schematic of *Act5C-RA* and *Act-RD* APA isoforms as well as their read coverage. Inset highlights the presence of the conserved ACACCC zipcode sequence present in the distal but not the proximal 3’ UTR.

Since RT terminations result when RNA is cross-linked to RBP, we mapped the location of 52 *Drosophila* RBP recognition motifs (29) defined by a Position Specific Scoring Matrix (PSSM) score (30). By clustering reads using the probability of an RBP motif within the 80-nt window from the RT termination site, we observed a subset of reads (15.1%) with an increase in the RBP motif probability (**Fig. 3d**). We named cDNAs termination sites with an RBP recognition motif as DORM (Downstream-Of-RBP-Motif). Out of 820 detected genes, we found 19 genes in the DORM cluster, including four ribosomal proteins (*RpL28, RpS16, RpS19a, RpS3A*), four cytoskeleton-associated proteins (*pod1, zip, Act5C, Dhc16F*), three proteins related to the Hippo pathway (*git, CG45050, GC42788*), *glec* (cell adhesion), *stau* (3’ UTR binding, A/P-axis specification), and *Zip48C* (solute carrier). For each gene, we calculated the ratio of cDNA reads terminating at or passing through predicted RBP motifs and ranked them by the frequency of DORM reads (**Fig. 3e**). We found that DORM cDNAs exhibited a shorter cDNA length (*t* = - 5.25, *p*-value < 0.0001) than non-DORM reads, containing more single-nucleotide mismatches at the 3’end (*OR*: 2.16, *CI*: 1.73 - 2.68, *p*-value < 0.0001). These properties were seen at the population level as well as at the level of individual genes (**Fig. 3h-i**). To identify *cis*-acting motifs at the DORM site in an unbiased manner, we clustered RBP motif *z*-scores from DORM reads (**Fig. 3j**) to identify probable RBP motifs (e.g. *msi, Sf1, Tra2, CG7804*) (**Fig. 3k**), which were color-coded for spatial analysis (**Fig. 3j**).

### INSTA-seq detects alternative splicing and polyadenylation (APA) isoforms

Tissue-specific alternative splicing or polyadenylation (APA) of the 3’ UTR is wide-spread (31). In addition, APA serves to generate 3’ UTR isoforms with an alternative set of *cis*-regulatory motifs affecting mRNA localization, degradation, or translation. However, spatially mapping APA isoforms has been challenging using *in situ* hybridization (ISH) due to a relatively small difference in their size, in addition to the abundance of repetitive or low-complexity sequences in the 3’ UTR (32).

To detect APA events in the *Drosophila* retina, we examined cDNA sequences near annotated polyadenylation sites. The nucleotide composition of the sequence flanking the last aligned position (LAP) was consistent with that of previously reported polyadenylation sites (33, 34) with U-rich Cleavage stimulation factor (CstF)-binding sites downstream and an A-rich polyadenylation signal (PAS) upstream (**Fig. 3l**). The CstF protein is necessary for the cleavage and polyadenylation, as well as for 3’ end processing of mRNAs. These results indicated that our *Drosophila* dataset was suitable for mapping sites of cleavage and polyadenylation (CPA). Subsequently, we identified ten genes that had at least two APA isoforms detected in the retina (*Act5C, Bacc, CG11360, Con, His4r, Nrg, Oda, ogre, Rtnl1, tws*), four of which are known to undergo tissue- or stage-specific extension of the 3’ UTR (35, 36) (**Fig. 3m**).

To see if APA changes the composition of regulatory elements in shorter 3’ UTRs compared to full-length 3’ UTRs, we examined proximal and distal 3’ UTR sequences flanking the CPA site. The median length for proximal 3’ UTRs was 385-nt (mean: 572-nt, s.d.: 441-nt) and 1,114-nt for distal 3’ UTRs (mean: 1,526-nt, s.d.: 1,177-nt). The median ratio of fragment sizes between proximal vs. distal 3’ UTRs was 46% (**Fig. 3n**), consistent with the hump observed across oligo(dT)-primed reads in the proximal 3’ UTR (**Fig. 3a**). Next, we performed motif analysis using *k*-mer-based differential enrichment for all proximal vs. distal 3’ UTRs pairs. The analysis revealed significant differential enrichment of C/A-rich sequences in the distal *Act5C* 3’ UTR (‘ACAC’, *p*-value < 0.01; ‘CACA’, *p*-value < 0.05) (**Fig. 3o**). The fraction of alternative isoforms ranged from 25% to over 95% across the ten genes, while *Act5C* expressed similar levels of short (*Act5C-RA*, 54%) and long isoforms (*Act5C-RD*, 46%) (**Fig.** 3p).

*Act5C* is one of six actin family genes in *Drosophila*, and its ortholog is the human cytoplasmic *β*-actin (*ACTB*). In *Drosophila*, the expression of *Act5C* is tissue- and developmental stage-specific (32, 37). The human *ACTB* mRNA is localized near the leading edge of migrating fibroblasts (38) as well as the growth cone (39), facilitating local translation and actin polymerization. The sub-cellular localization of the *ACTB* mRNA is mediated by the IGF2-mRNA binding protein (IMP1/ZBP1) (40, 41). IMPs are a family of homologous RBPs that are conserved from insects to mammals (42). Studies have examined IMP1-binding motifs using cross-linking and immunoprecipitation (CLIP), and such assays have identified C/A-rich sequences, as well as CA dinucleotide motifs recognized by the third KH domain of IMPs (43). The IMP1 *Drosophila* ortholog dIMP binds two A/C-ACA motifs separated by 0-30 nucleotides (44), which were detected in the distal *Act5C* 3’ UTR but not in the short *Act5C* isoform (**Fig. 3q**). Since dIMP knockout leads to multiple defects in cell migration and growth cone dynamics (44), it implies that APA could regulate *Act5C* mRNA localization through ZIP/IMP *cis*-acting motifs during neuronal development (45).

### INSTA-seq maps transcripts, regulatory motifs, and APA tissue-wide with subcellular resolution

INSTA-seq is compatible with immunohistochemistry or fluorescent proteins, enabling one to examine localization patterns relative to the protein of interest. In our case, we imaged *α*-catenin-GFP (**Fig. 4a**) prior to *in situ* sequencing, and we subsequently applied a deep Conditional Adversarial Network (46) (**Fig. S4**) to automatically segment cell-cell boundaries of the three major cell-types that define the ommatidium morphology. This enabled us to assign individual cDNA amplicons within each individually segmented cell (**Fig. 4b-c**). Cell-cell boundaries in an individual ommatidium was then used to automatically register individual cDNA amplicons onto a subcellular geometric reference atlas coordinate system (**Fig. 4d-e**).

**Fig. 4.**
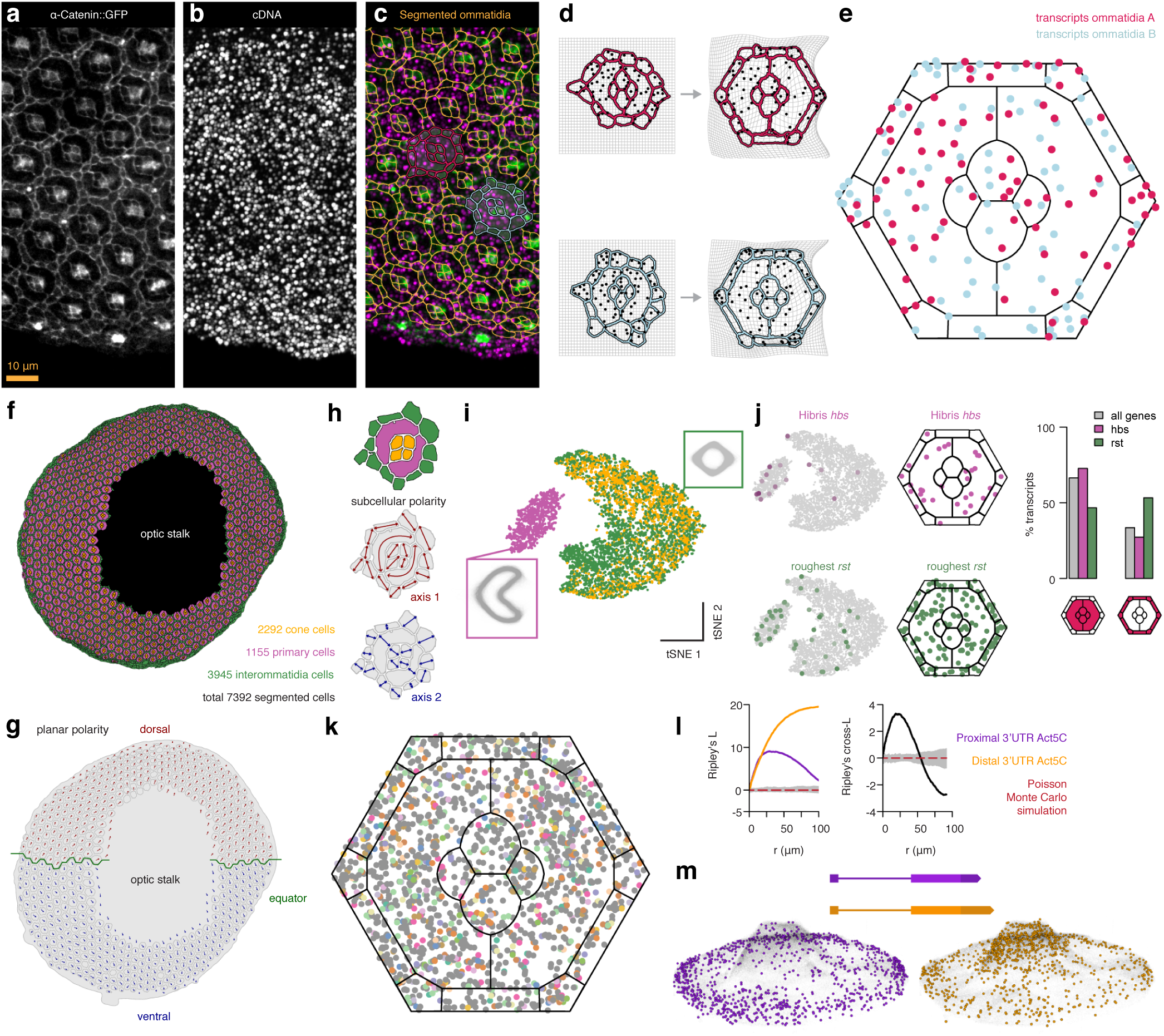
Subcellular and tissue-wide localized gene expression. **a)** *α*-catenin::GFP expression in the *Drosophila* retina. **a)** cDNA amplicons hybridized with fluorescent sequencing primer. **c)** Segmented ommatidium (orange) overlaid on cDNA amplicons (magenta) and *α*-catenin::GFP (green). **d)** Two individual ommatidias highlighted in c) with segmented ( in situ) amplicons transformed into the ommatidium reference atlas. (e) Amplicons from ommatidia from d) visualized in the common coordinate reference atlas. **f)** 7392 single cell boundaries segmented from a single *Drosophila* retina. **g)** Planar polarity assigned to each ommatidia (arrows) as well as the equator (green). **h)** Cellular polarity extracted for each cell contour. **i)** Multidimensional reduction of cell morphology. Insets shows cell contours superimposed on top of each other each contour drawn using transparent gray color. **j)** *Hbs* and *rst* gene expression superimposed on a multidimensional representation of cell morphology. Individual amplicons of *hbs* (magenta) and *rst* (green) superimposed on the ommatidium reference atlas. Bargraph of the percentage of amplicons in IOCs vs. cone or primary pigment cells. **k)** Amplicons from DORM reads in Fig. 2j clusters visualized within the reference ommatidium. **l)** Left: Ripley’s *L* as a function of radius in micrometers for proximal (purple) and distal (orange) isoforms. Right: Cross Ripley’s *L* between proximal and distal isoforms. **m)** Spatial distribution in entire *Drosophila* retina of *Act5C* proximal (purple) and distal (orange) single transcripts.

In total, 774 ommatidium boundaries were segmented, except near the optical stalk where the GFP signal was low. 573 ommatidia were segmented into 2292 cones cells, 1155 primary cells, and 3945 inter-ommatidia cells (IOC) (**Fig. 4f**). We used the GFP signal in the apical planes to assign planar polarity to each ommatidia as well as tissue-wide developmental axes (dorsal-ventral) and the equator (**Fig. 4g**). In addition, we used individual cell morphology to compute the first two principal components of the cell contour and generated a two-dimensional subcellular polarity coordinate (**Fig. 4h**). Multidimensional reduction of cell contours provides an unbiased way to extract average cell morphologies and clusters (**Fig. 4i**). We validated our computational framework by examining known cell type-specific markers. The adhesion proteins Roughest (*Rst*) and Hibris (*Hbs*) control cellular organization through cell sorting by differential adhesion (47). *Hbs* is preferentially expressed in primary pigment cells, while *rst* is preferentially expressed in IOCs. Preferential Hbs-Hbs interactions between 1° pigment cells, and Hbs-Rst interactions between 1° cells and IOCs contribute to sorting of 1° cells from IOCs. Although our pilot dataset did not have deep sequencing coverage (**Fig. 4j**), the relative expression levels of *hsb* and *rst* was consistent with the reported protein expression (OR: 0.33, CI: 0.13 - 0.80, p-value 0.008; **Fig. 4j**). At the current sequencing depth statistically significant polarization of *cis* motifs or 3’ UTR-RBP interactions was not found (**Fig. 4k**); however, we expect more sequencing data to reveal asymmetrically localized regulatory sequences and interactions.

To define the expression patterns of the short and long *Act5C* isoforms, we first performed an unbiased point pattern analysis. We found that the proximal isoform aggregated at a spatial scale of 30 µm, while the distal isoform aggregated on larger spatial scales (**Fig. 4l**). At smaller spatial scales (<50 µm) the two isoforms tended to co-cluster, while on larger spatial scales the two isoforms dispersed away from each other as shown by a negative cross Ripley’s *L* function (**Fig. 4l**) that ask whether any two point co-localization patterns deviate from random probability distributions. On closer examination this dispersion was due to higher aggregation of the distal isoform in the optic stalk of the retina (*OR*:9.52, *CI*:7.53-12.10, *p*-value < 0.0001), **Fig. 4m**), a region known for supporting retinal glial cells migrating from the optic stalk into the eye disc (48). This is consistent with the presence of a zipcode consensus motif in the distal 3’ UTR and subsequent localization and translation of the actin mRNA necessary for cell migration (49).

## Discussion

Cell type- or context-dependent interactions between RNA, and proteins are essential to identifying functional genetic elements and deciphering their function and modes of regulation. To do so, we developed INSTA-seq for mapping mRNA-RBP interactions with subcellular resolution *in situ*. The INSTA-seq method is separated into three parts: 1) UMI sequencing *in situ* using PRICKLi for scalable imaging, 2) mated-pair NGS for longer reads and 3) spatial mapping of mRNAs with subcellular resolution. Notably, the sequencing chemistry we developed can be customized to a wide range of applications beyond INSTA-seq. Consistent with the previous observations obtained using FISSEQ (12), we discovered that INSTA-seq detects tissue- or pathway-specific gene clusters despite utilizing a minimal number of sequencing reads. It also contains cross-linking signatures for mapping 3’ UTR-RBP interactions across the tissue *in situ*. By placing sequencing reads into morphology-based reference bins, locally enriched 3’ UTR-RBP interactions can be identified with subcellular resolution with more reads. In addition to mRNA-protein binding, we found that alternative polyadenylation (APA) alters zipcode binding protein motifs in the *Act5C* 3’ UTR, revealing a functional consequence of tissue-specific APA (45). Finally, tissue- or pathway-specific transcripts are selectively amplified while abundant ribosomal protein mRNAs escape detection. We hypothesize that a subset of the transcriptome is more open to dynamic regulation, affecting their accessibility to reagents specific to INSTA-seq. We are now investigating RNA-protein interaction and RNA localization patterns across cell types and perturbations to identify the source of detection bias.

## ACKNOWLEDGEMENTS

We would like to acknowledge the following individuals for their help T. Chan (MSKCC) for providing cell lines, O. Chaudhary (CSHL), B. Bossman and N. Claxton (Nikon) for microscopy setup, D. Ghosh (CSHL) and K. Lao (ThermoFisher) for SOLiD sequencing, M. Hammell (CSHL), and K. Mir (XGenomes). We would also like to thank former and present members of Lee, Wigler, Hammell, and Hicks labs for their assistance, as well as CSHL sequencing and computing facilities. Finally, we would like to thank A. Kepec for helpful suggestions on preparing the manuscript. This work was supported by NIGMS R35 RFA-GM-003, The V Foundation, STARR Cancer Consortium, CSHL-Northwell translational grant, and Pershing Square Scholarship to J.L. and by the National Institutes of Health NIGMS GM129151 to V.H.

## METHODS

### PRICKLi probes

All oligos were synthesized by Integrated DNA Technologies, Inc. (IDT). Full design and ordering specifications are outlined in **Table S3-6**. Reverse set PRICKLi probes due to their single ribonuclotide were ordered as custom RNA oligos. All other oligonucleotides were ordered as custom DNA oligos.

### Tissue dissection and mounting

*Drosophila* retinas expressing UAS-*α*-Catenin∷GFP with the eye-specific GMR-GAL4 driver (50) for highlighting cell outlines were dissected at 26 to 28 hours after puparium formation (APF). Retinas were then fixed, washed in phosphate buffer, and mounted with the retina apical surface facing an acid-washed cover glass coated with Bio-Bond tissue adhesive (Ted Pella Inc.). Each cover glass contained 9 wells and several retinas were mounted per well. The retinas were partially dried on the cover glass to facilitate tight boding with the glass surface. Then, the coverslips were fitted with an adhesive silicon isolator (Grace Bio-Labs) to simplify subsequent liquid handling and tissue imaging.

### Cell culture

Immortalized human astrocytes (51) were grown in Dulbecco’s Modified Eagle’s Medium (DMEM) plus 10% fetal bovine serum (FBS; Invitrogen) on fibronectin-coated (F1141, SigmaAldrich) plates. WA09 FUCCI hESC (52) were cultured in mTeSR1 (85850, STEM-CELL Technologies) with with 1 × RevitaCell ROCK inhibitor (A2644501, ThermoFisher) and Matrigel Matrix (EMD Millipore).

### *In situ* library preparation

Frozen retinas were thawed and washed in 1x PBS in DEPC H2O (750023, ThermoFisher). Post-fixation was done in 10% formalin (HT5011-1CS, Sigma-Aldrich) for 15 min followed by three washes in 1xPBS in DEPC H2O. Tissues were permeabilized in 0.3% TritonX-100 (93443, Sigma-Aldrich) in DEPC H2O with 0.5 U/µl of SUPERase (AM2696, ThermoFisher Inc) for one hour. After permeabilization the nuclear envelope was permeabilized with 0.1 HCl in DEPC H2O for 45 min followed by three washes in 1xPBS in DEPC H2O.

Genomic DNA (gDNA) was digested by 5 µl of TURBO DNase (AM1907, ThermoFisher) in 175 µl nuclease-free water with 20 µL of 10 × Turbo DNA-free buffer (AM1907, ThermoFisher) incubated at 37° C for one hour. After gDNA digestion it is important to inactivate DNase I with by resuspending with DNase Inactivation Reagent (0.2 of the volume) into the reaction mix. Incubate for 30 min at room temperature on a shaker. The sample was rinsed with ice cold ddH2O several times. The inactivation reagent leaves white precipitate that was thoroughly wahsed away using nuclease-free water.

Next, primer hybridization was done with 2.5 µM of anchored oligo(dT)10 INSTA-seq reverse transcription primer in DEPC-2xSSC with 0.5 U/µl of SUPERase (AM2696, ThermoFisher Inc) overnight at room temperature. In the morning the sample was placed on ice, and the reverse transcription buffer reaction mix without enzymes was prepared on ice [in DEPC-H2O, 1 × M-MuLV RT buffer (P7040L, Enzymatics), 250 µM dNTP, 25 mM (N2050L, Enzymatics), 40 µM Aminoallyl-dUTP, 4 mM (83203, AnaSpec)]. The primer was aspirated on ice, and the sample was pre-incubated on ice with the RT reaction mix without any enzyme for 15 min. The sample was aspirated on ice, and the revere transcription reaction mix was added with enzymes (same as above but with 5 U/µl of M-MuLV [P7040L, Enzymatics], 0.5 U/µl of SUPERase). The sample was incubated for 15 min on ice, followed by 15 min at 25° C and then at 37° C overnight.

The following day the sample was gently aspirated and 20 µl of BS(PEG)9 (21582, ThermoFisher) dissolved in 980 µl of 1xPBS pH 8 (CHP-300, Boston BioProducts), which was then added to the sample to cross-link newly synthesized cDNA for one hour. BS(PEG)9 was then aspirated and quenched by incubating in 1 M Tris-HCl pH 8.0 (AM9855G, ThermoFisher) for 30 min. To remove RNA in the RNA:cDNA duplex the sample was incubated for one hour at 37° C with 125 mU/µl RNase H (Y9220F, Enzymatics) and 500 ng/µl RNase A (11579681001, Roche) in 1 × Rnase H buffer (Y9220F, Enzymatics). After RNA removal the sample was thoroughly rinsed with nuclease-free water three times. The sample was placed on ice and preincubated with the CircLigase II reaction buffer on ice for 5 min (Nuclease-free H2O, 1 × CircLigase buffer, 2.5 mM MnCl_2_, 0.5 M Betaine, [all included in kit from CL9025K, Epicentre]). After aspiration the CircLigase II reaction mix (same buffer composition as previous but with 1U/µl of CircLigase II enzyme, CL9025K, Epicentre) was added to the sample and incubated for two hours at 60° C. The sample was aspirated and prehybridized with 1 µM oligodT10 exonuclease protected RCA primer in 2xSSC with 30% (vol/vol) formamide (221198, SigmaAldrich) at 60° C for two hours. The sample was then aspirated and incubated first for 15 min at 60° C with 10% (vol/vol) formamide in 2xSSC, followed by 2xSSC for another 15 min at 60° C. The sample was allowed to cool to room temperature and then placed on ice. The sample was preincubated with the RCA reaction mix without enzyme on ice for 5 min, followed by the RCA reaction mix with high concentration *ϕ*29 polymerase (1 × *ϕ*29 buffer, 250 µM dNTPs, 40 µM Aminoallyl-dUTP, 1 U/µl high-concentration *ϕ*29 DNA polymerase, P7020-HC-L, Enzymatics) incubated at 30° C for no less than 16 hours. After RCA the sample was gently aspirated, and BSPEG9 dissolved in 1xPBS pH 8 was used to fixate rolling circle products (RCPs) for one hour. This is followed by aspiration and quenching with 1 M Tris-HCl pH 8.0 for 30 min, followed by a wash in 1xPBS. After the fixation *in situ* cDNA library quality is ready to be assessed by hybridizing a fluorescent sequencing primer onto the RCPs by heating up 2.5 µM of fluorescent sequencing primer in 5xSASC (0.75 M sodium acetate, 75 mM tri-sodium citrate, adjusted pH to 7.5 using acetic acid) to 80° C and by adding the primer solution onto the sample. After cooling to room temperature for 15 min, several washes are performed with 5xSASC, and then the sample is ready to be imaged. The RCP density, roundness, brightness and overall sample quality are examined before *in situ* sequencing. Endogenous fluorescent proteins are imaged at this step to ensure best signal to noise before any ligation or cleaving of fluorescent sequencing probes is performed.

### *In situ* sequencing

In SBL sequencing is partitioned into ligation cycles and hybridization rounds. In ligation cycles the sequencing primer is extended by ligation. In hybridization rounds a new primer (usually a recessed version of a primer from previous primer round) is hybridized and then extended for a predetermined set of ligation cycles until the entire extended primer is stripped away and a new round is started. PRICKLi uses three primer rounds and two ligation cycles in each direction for each primer round. It is important that forward probes are ligated before reverse probes.

First, any remaining fluorescent sequencing primer is stripped away with stripping buffer 80% (vol/vol) formamide (221198, Sigma-Aldrich) in H2O and 0.01% (vol/vol) Triton X-100 pre-heated to 80° C and incubated for 5 min twice with three washes of constant flow of 10 ml 5xSASC thereafter. Sequencing primers (2.5 µM) are heated to 80° C in 5xSASC and directly applied onto the tissue to anneal at room temperature for 15 min followed by three washes in 5xSASC afterwards. Ligation of forward probes is then done with 2.5 µM of equimolar forward probes in 1 × T4 DNA ligase buffer and 6 U/µl T4 DNA ligase (L6030-LC-L, Enzymatics) for minimum of 45 minutes, followed by three washes of constant flow of 10 ml 5xSASC. Reverse probes are ligated the same way followed by three washes of constant flow of 10 ml 5xSASC. The sample is then ready to be imaged for the first round. After imaging, the probes are cleaved to enable a second ligation cycle.

Cleaving is done with 0.2 U/µl of Endonuclease V (m0305, NEB) and 0.2 U/µl of RNase H2 (M0288L, NEB) in 1 × NEBuffer 4 (both enzymes in the same reaction). Fluorophores are cleaved within 5 minutes, but it is advised to let the reaction run to completion of 45 min to minimize dephasing. The sample is washed extensively with a constant flow of 10 ml 5xSASC afterwards three times. The sample is now ready for another ligation cycle.

After two ligation cycles, the extended sequencing primer is cleaved and stripped away with stripping buffer 80% (vol/vol) formamide (221198, Sigma-Aldrich) in H2O and 0.01% (vol/vol) Triton X-100 pre-heated to 80° C and incubated for 5 min two times with three washes using a constant flow of 10 ml 5xSASC. A new set of sequencing primers (2.5 µM in 5xSASC) are then hybridized by heating primers to 80° C and directly applying them onto the tissue and letting them cool to room temperature for 15 min, followed by three washes in 5xSASC.

### Image acquisition

*In situ* sequencing was performed on a Nikon Ti-E Inverted Microscope with PFS3 Yokogawa CSU-X1 spinning disc confocal (Nikon). FOV pixel size of 1600 × 2048 px and a theoretical pixel resolution of 0.11 µm using a 40 × NA 1.1 CFI Lambda S Apo LWD 4 Water Immersion Objective Lens (MRD77410, Nikon) (9-12 tiles per retina, 90-200 *z*-stacks with a 500-nm step size). With LU-NV NIDAQ lasers (488 nm, 561 nm, 594 nm, 640 nm) with 8900 Sedat Quad dichroic mirrors (425-477, 503-542, 571-628 and 661-728 nm).

### *In situ* base calling

Raw TIFF image stacks were first corrected for chromatic shift (53). They were then flat-field corrected and segmented using wavelet filters from *Whole-Brain* R package (54). After segmentation of 3D connected components across *z*-stacks, the overlap between channels for individual amplicons was calculated using Manders co-occurrence (55) between all pixels in 3D. The co-occurrence index was then weighted by the fluorescent intensity using probit regression, and the final score was used to cluster individual amplicons into single base calls using *K*-means clustering in R.

### *In situ* registration

At each imaging cycle, 3D amplicon centroids in all channels were registered back onto previous imaging cycle using the Coherent Point Drift equation (56) implemented in C++. The resulting correspondence index vector was used to match back amplicons across imaging cycles. Amplicons that disappeared and reappeared across imaging cycles were designated as cytosine-cytosine base calls for the image cycle where they disappeared. The auto-correlation function across all amplicons and cycles was used to subtract any remaining dephasing and base calls were reclustered by *K*-means clustering. The final set of base call reads for individual amplicons in addition to the quality metrics of base calling based on cluster separation was written to paired-end R1 and R2 FASTQ files.

### NGS library preparation

After *in situ* sequencing the amplicon quality was inspected visually, and any fluorescent probes were stripped and thoroughly washed away. The tissue quality was inspected under bright-field microscopy. Since BSPEG9 fixation makes the tissue hardened, a pair of small forceps (11255-20, Fine Science Tools) was used to loosen the tissue from the coverslip. Next, the tissue was placed into an Eppendorf tube with 20 µl of lysis buffer (200 mM KOH, 40 mM DTT, 5 mM EDTA). The Eppendorf tube was placed in a 95° C heat block for 10-20 minutes until the tissue was determined to be completely lysed by visual inspection. For cell cultures, the lysis buffer was added directly to the well and pipetted up and down, and the pipette tip was used to scratch away cells from the glass bottom. The lysis buffer and cells were added to the Eppendorf tube and placed on a heat block. The well plate was inspected under bright-field to make sure all cells were removed. After lysis the tube was vortexed, and 20 µl of neutralization buffer (400 mM HCL, 600 mM Tris-HCl pH7.5) was added to the tube, followed by 10-minute incubation on ice. When the neutralization buffer has been added, the entire 40 µl solution can be stored at −80° C until further processing.

Next, 250 µl Dynabead M-270 Streptavidin (65306, ThermoFisher Inc) was washed on a magnetic stand three times in 2xSSC. Biotinylated INSTA-seq pulldown oligo was thawed and diluted in 2xSSC (2.5 µM, 5 µl in 195 µl 2xSSC). Dynabead was placed on the magnetic stand for 5 min, and 2xSSC was aspirated on the stand. The pulldown oligo in 2xSSC was added to the beads and placed on a shaker for 15 min. The tube was then placed on the magnetic stand for 5 min, and any 2xSSC supernatant was aspirated on the stand. Immediately, the 40 µl solution containing the lysed *in situ* tissue was added to the beads and filled to 200 µl with 2xSSC. The tube was placed on an 80° C heat block for 5 min and then was placed to cool down at room temperature for 15 min and then placed on ice. The tube was placed on the magnetic stand for 5 min, and the supernatant was aspirated on the magnetic stand and washed with 2xSSC two times on the magnetic stand.

Next, second strand synthesis was performed by removing any supernatant on the magnetic stand and by immediately adding the ice cold phi29 second strand synthesis reaction mix (176 µl of nuclease-free water, 20 µl phi29 10x buffer, 2 µl of 25 mM dNTPs, 2 µl of phi29 DNA polymerase [100 U µl-1]) and placed in a 30° C incubator overnight.

The following day the tube was placed on the magnet for 5 min, and the supernatant was aspirated and discarded. While on the magnet, 10 µl of 2x annealing buffer (10 mM Tris pH8, 50 mM NaCl, 1 mM EDTA) was added to the beads and gently pipetted up and down several times. Exonuclease-resistant random hexamer primers 1 µl (Thermo SO181) was added into the reaction mix and the beads were suspended in nuclease-free water to 20 µl. The tube with the beads, exonuclease-resistant random hexamers, and annealing buffer was placed on a 95° C heatblock for 5 min, followed by cooling down on ice to 4° C degrees. While on ice, the multiple displacement amplification (MDA) reaction mixture was made by supplementing the 20 µl bead solution with 5 µl of 10x phi29 DNA polymerase reaction buffer, 2 µl of 25 mM dNTPs, 21 µl of nuclease-free water, and 2 µl of phi29. The tube was placed in a 30° C incubator, and one microliter was withdrawn from the sample after three hours to check the dsDNA concentration using Qubit dsDNA HS Assay Kit (Thermo Q33230) on a Qubit 4 Fluorometer (Thermo Q33238). If amplification was successful, the sample was further amplified at 30° C for 10 hours, followed by a 3-minute incubation at 65° C to inactivate the phi29 polymerase.

The dsDNA concentration of the final sample was measured again using Qubit 4 Fluorometer, and the Nextera DNA FLEX kit (20018704, Illumina) libraries were prepared. Briefly, the dsDNA was tagmented by bead-linked transposomes followed by clean up. Index adapters (FC-131-2002, Illumina) were added to multiplex different tissues or samples, and tagmented DNA was amplified using five cycles of PCR according to the manufacturer’s instructions. The amplified libraries were double-sided bead purified, followed by a quality control step using 2100 Bioanalyzer with a High Sensitivity DNA kit (Agilent). The samples were pooled, diluted, and prepared according to the MiSeq kit specification (either v3 or v2). We evaluated both 75-bp and 300-bp read length, and the benefit of 300-bp is observed for cDNA lengths greater than 85 nt.

### Alignment

Reads containing the *in situ* sequencing adapter were split using removesmartbell.sh from BBmap (57). Any adapters were trimmed, and paired-end reads were aligned to the *Drosophila* melanogaster genome version BDGP6 (Ensembl) using BBMap (57), HISAT2 (58) or STAR (59). For comparison to published single-cell RNA-seq data (27), STAR was used as aligner. Multiple cDNA reads mapping to the same genomic coordinate with the same UMI were assigned to the same *in situ* amplicon. Individual *in situ* amplicons were assigned genomic features using featureCounts (60).

### DORM reads clustering

The cDNA termination site was assigned to each read, and 51 RNA binding protein motifs from the RBPmap full list iand dIMP (44) were mapped to the 3’ UTR for each gene using RBPmap (30). The bed-tracks of *z*-score prediction summaries for each RBP recognition motif in a window of 80-nt near the cDNA termination site was used as input for hierarchical clustering. Euclidean dissimilarities were calculated for the resulting matrix, and the function hclust with Ward method from R was used to hierarchical cluster the reads.

### Segmentation of cell boundaries using *α*-catenin with generative adversarial networks

One and a half retina were manually annotated with individual cell boundaries drawn based on *α*-catenin GFP signal. Regions of interests were manually drawn in ImageJ, binarized and saved as 256×256 tiles together with bandpassed filtered raw *α*-catenin-GFP signal as a training set to pytorch implementation of pix2pix (46).

### Registration of ommatidia and amplicons to the reference atlas ommatidium

Pixels of segmented cell boundaries from *α*-catenin∷GFP were registered to pixels of a standardized reference atlas ommatidium using the Coherent Point Drift equation (56) implemented in C++. The resulting transformation field was then used to transform the location of individual *in situ* amplicons to the reference atlas. Before registration the planar polarity of each ommatidium was taken into consideration by simple reflection such that both dorsal and ventral ommatidia share the same chirality in the reference atlas.

### Multidimensional reduction of cellular morphology

Cell boundaries were segmented based on previous described adversial network (46) on *α*-catenin-EGFP signal. Pixel coordinates for each individual contour was first reduced to 200 pixels with spsample using the sp library in R. Next, the first two principal components of each contour were computed and the intersection of the first principal component was used as zero-index of the contour. For primary pigments cells, the first principal component was replaced by the medial axis. Both *x* and *y* coordinates were appended into a 400 dimensional matrix where each row represents individual cell boundaries and the first 200 columns represents *x*-values from the contour and the last 200 columns represents *y*-values from the contour. Both *x* and *y* values were centered by the intersection of the first two principal components and then scaled by the standard deviation. The resulting 7242 × 400 array was then used as an input into the t-SNE algorithm using Rtsne package in R.

### *In vitro* optimization

In vitro reactions of ligation and cleaving was optimized with fluorescent oligos using readout with electrophoresis of denatured ligated or cleaved product on 15% TBE-urea polyacrylamide gel and visualized by Typhoon FLA 7000 imager (GE Lifesciences).

### Data analysis

All analysis and plotting was done in R (3.5.3) with statistical tests including Fisher’s exact test, chi-square goodness of fit, student *t*-test and linear regression. For spatial analysis Ripley’s *L* was calculated using the spat-stat R package.

---

**Supplementary Figure 1.**
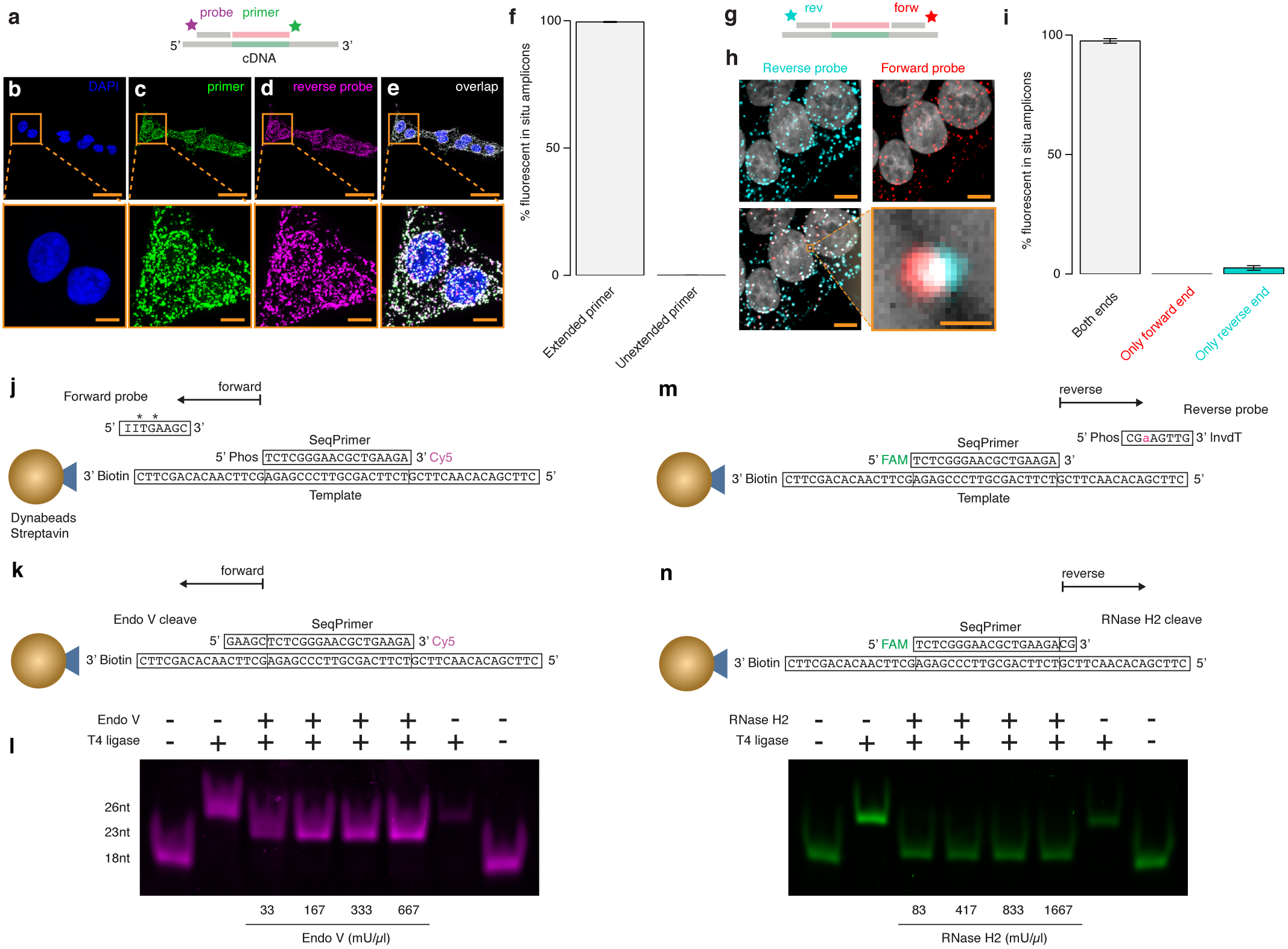
Characterization of PRICKLi fluorescence ligation, co-localization, and cleavage *in vitro*. **a)** *In situ* ligation efficacy in the reverse direction was evaluated on cDNA amplicons in human embryonic stem cells. Amplicons were hybridized with a 5’ FAM-labeled sequencing primer (18-nt), followed by ligation of Cy5-labeled complete degenerate reverse PRICKLi probes using T4 DNA ligase for 45 min (scale bar: 2-µm). **b)** DAPI staining. **c)** FITC fluorescence. **d)** Cy5 fluorescence. **e)** Co-localization of FAM and Cy5 fluorescence. **f)** Percentage of FAM-labeled sequencing primers on cDNA amplicons extended by PRICKLi. **g)** cDNA amplicons hybridized to the unlabeled sequencing primer, followed by sequential ligation of complete degenerate Cy3-labeled forward PRICKLi and complete degenerate Cy5-labeled reverse PRICKLi probes. **h)** Sparse amplicon generation by shortening cDNA synthesis incubation makes co-localization of Cy3 and Cy5 fluorescence unambiguous by visual inspection. **i)** Percentage of sequencing primers on cDNA amplicons extended by forward and reverse PRICKLi probes. **j)** Oligonucleotide design for testing forward PRICKLi ligation and textbfk) cleavage. The ligated product was denatured and analyzed on a 15% TBE-Urea gel, demonstrating complete ligation (26-nt) using T4 DNA ligase after 45 min and complete Endonuclease V-dependent cleavage (23-nt) after 45 min. **m)** Oligonucleotide design for testing reverse PRICKLi ligation and cleavage. **n)** The ligated product was denatured and analyzed on a 15% TBE-Urea gel, demonstrating complete ligation (26-nt) using T4 DNA ligase after 45 min and complete RNase H2-dependent cleavage (20-nt) after 45 min.

**Supplementary Figure 2.**
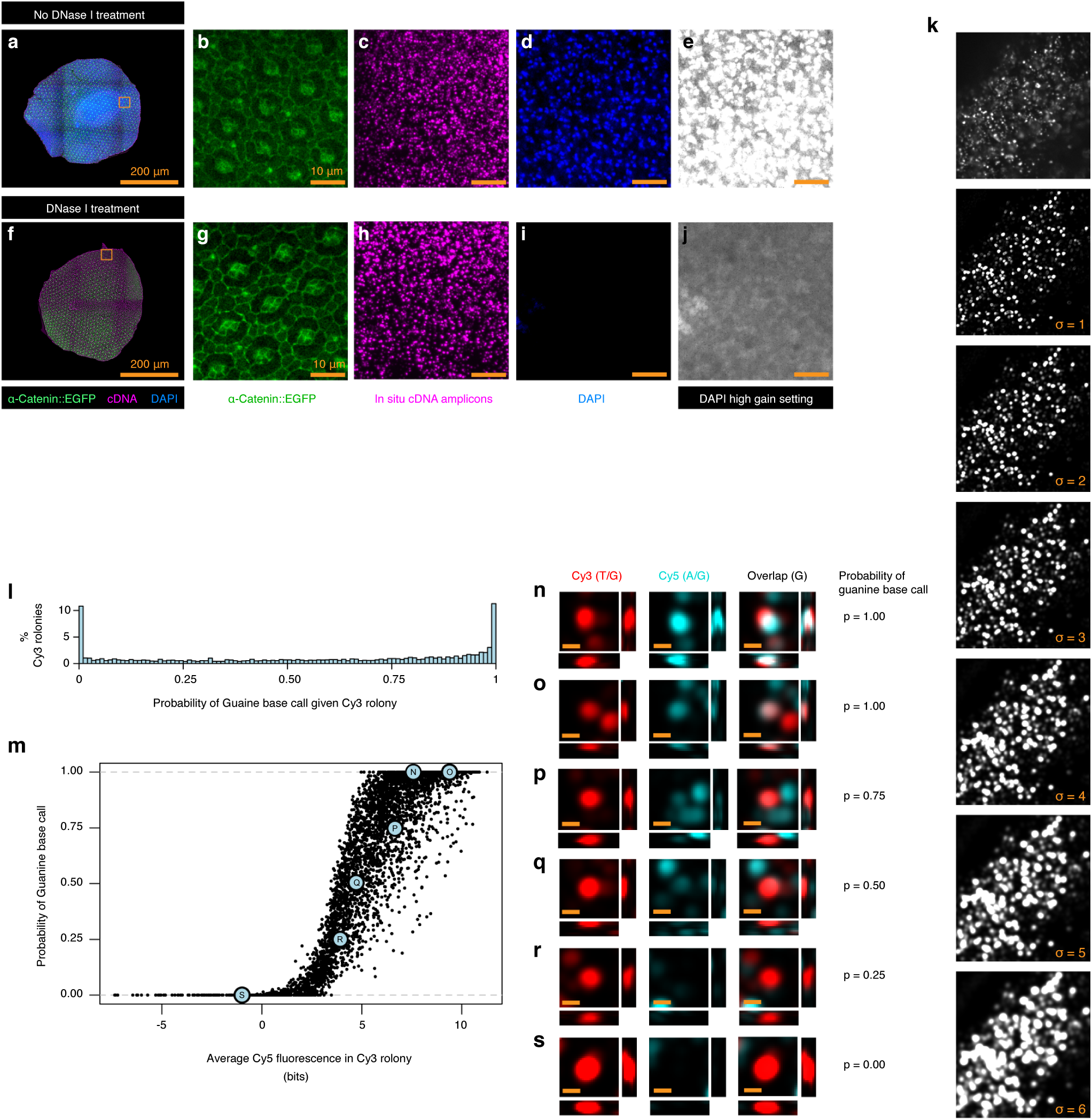
Tissue and image processing of INSTA-seq samples. Genomic DNA digestion using DNase I does not affect subsequent *in situ* amplicon cDNA quality or density. **a)** DAPI stained *Drosophila* retina hybridized to the Cy5-labeled sequencing primer, close up in **b-e** seen as orange square. **b)** *α*-catenin-GFP signal. **c)** Cy3-labeled cDNA amplicons. **d)** DAPI staining of the genomic DNA. **e)** DAPI staining of the genomic DNA (high gain). **f)** DAPI stained *Drosophila* retina (post DNase I-treatment) hybridized to the Cy5-labeled sequencing primer. g) *α*-catenin-GFP signal. **h)** Cy3-labeled cDNA amplicons. **i)** DAPI staining of the genomic DNA. **j)** DAPI staining (high gain). **k)** Top shows raw fluorescent image from single *z*-plane slice from a single channel during *in situ* sequencing. Subsequent images at the bottom shows wavelet processed output where energy at a scale period of roughly the size of a single *in situ* amplicon is used as input to compute tensor structure and to extract trace energy with different Scharr operator kernel sizes for enlarging amplicons. This step makes *in situ* amplicons more uniform and improves co-localization analysis. Typically a kernel size of *σ* = 2 is used for downstream analysis. **l)** Manders co-occurance index (55) between two channels generates a conditional probability measurement of the probability of guanine base call, given that the amplicon was originally segmented using other color channels. The probability measurement roughly follows Jeffrey’s prior for a Binomial likelihood function, ∼ *Beta*(1/2, 1/2). **m)** Probability of a guanine base call for Cy3 segmented rolonies as a function of the average Cy5 fluorescent signal. Individual amplicons highlighted in light blue are also shown in **n-s**, scale bar: 1 µm.

**Supplementary Figure 4.**
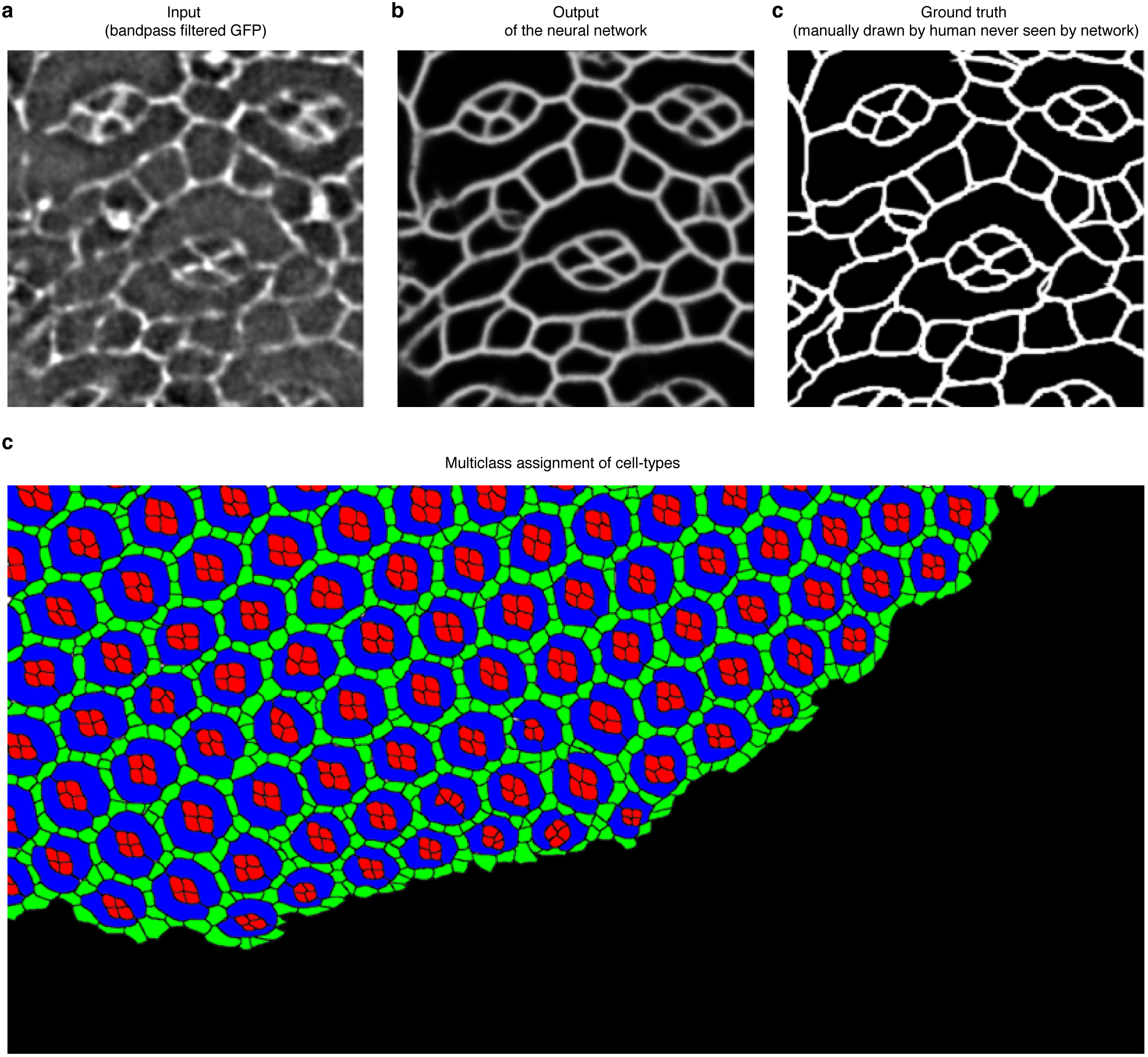
Segmentation of ommatidia with Conditional Adversarial Networks. **a)** *α*-catenin-GFP signal was first band-pass filtered and provided as input. **b)** Output of trained network given input tile from **a). c)** Ground truth. **c)** Multi-class assignment of individual cell-types into three different channels. Red: cone cells, Blue: primary pigment cells, Green: inter-ommatidia cells.

**Table 1.**
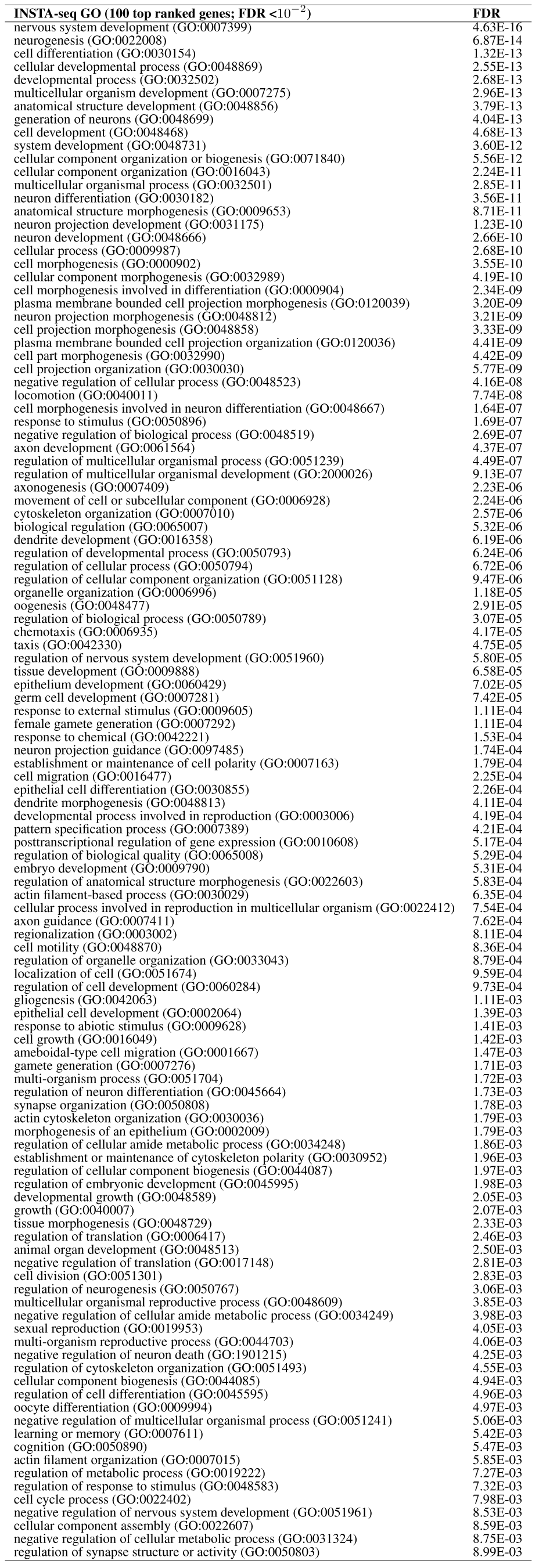
INSTA-seq GO term enrichment in the *Drosophila* retina.

**Table 2.** DROP-seq GO term enrichment in the *Drosophila* eye imaginal disc (27)

| DROP-seq GO (100 top ranked genes; FDR <10<sup>-2</sup>) | FDR |
| --- | --- |
| cytoplasmic translation (GO:0002181) | 5.57E-122 |
| translation (GO:0006412) | 2.75E-100 |
| peptide biosynthetic process (GO:0043043) | 5.51E-100 |
| amide biosynthetic process (GO:0043604) | 1.49E-97 |
| peptide metabolic process (GO:0006518) | 6.56E-93 |
| cellular amide metabolic process (GO:0043603) | 3.25E-87 |
| cellular nitrogen compound biosynthetic process (GO:0044271) | 3.32E-73 |
| organonitrogen compound biosynthetic process (GO:1901566) | 6.81E-73 |
| cellular macromolecule biosynthetic process (GO:0034645) | 2.24E-72 |
| macromolecule biosynthetic process (GO:0009059) | 1.64E-71 |
| gene expression (GO:0010467) | 5.26E-65 |
| cellular biosynthetic process (GO:0044249) | 3.43E-57 |
| organic substance biosynthetic process (GO:1901576) | 5.27E-56 |
| biosynthetic process (GO:0009058) | 2.51E-55 |
| cellular protein metabolic process (GO:0044267) | 1.63E-50 |
| cellular nitrogen compound metabolic process (GO:0034641) | 5.54E-50 |
| cellular macromolecule metabolic process (GO:0044260) | 1.22E-42 |
| protein metabolic process (GO:0019538) | 2.91E-42 |
| organonitrogen compound metabolic process (GO:1901564) | 1.91E-35 |
| macromolecule metabolic process (GO:0043170) | 1.15E-31 |
| nitrogen compound metabolic process (GO:0006807) | 1.86E-28 |
| cellular metabolic process (GO:0044237) | 1.32E-26 |
| primary metabolic process (GO:0044238) | 1.78E-26 |
| ribosome biogenesis (GO:0042254) | 6.80E-24 |
| organic substance metabolic process (GO:0071704) | 8.52E-24 |
| cellular process (GO:0009987) | 5.16E-22 |
| metabolic process (GO:0008152) | 5.83E-21 |
| ribonucleoprotein complex biogenesis (GO:0022613) | 2.20E-20 |
| ribosome assembly (GO:0042255) | 7.29E-18 |
| ribosomal large subunit biogenesis (GO:0042273) | 1.65E-16 |
| ribonucleoprotein complex assembly (GO:0022618) | 2.49E-12 |
| ribonucleoprotein complex subunit organization (GO:0071826) | 7.18E-12 |
| ribosomal small subunit biogenesis (GO:0042274) | 4.08E-11 |
| cellular component biogenesis (GO:0044085) | 1.24E-09 |
| ribosomal large subunit assembly (GO:0000027) | 2.64E-09 |
| cellular protein-containing complex assembly (GO:0034622) | 9.78E-09 |
| ribosomal small subunit assembly (GO:0000028) | 1.08E-08 |
| protein-containing complex assembly (GO:0065003) | 5.51E-08 |
| rRNA processing (GO:0006364) | 6.09E-07 |
| Unclassified (UNCLASSIFIED) | 6.50E-07 |
| biological_process (GO:0008150) | 6.66E-07 |
| rRNA metabolic process (GO:0016072) | 8.89E-07 |
| organelle assembly (GO:0070925) | 1.52E-06 |
| protein-containing complex subunit organization (GO:0043933) | 1.60E-06 |
| maturation of SSU-rRNA (GO:0030490) | 2.43E-04 |
| ncRNA processing (GO:0034470) | 2.64E-04 |
| cellular component assembly (GO:0022607) | 8.40E-04 |
| maturation of SSU-rRNA from tricistronic rRNA transcript (GO:0000462) | 1.17E-03 |
| ncRNA metabolic process (GO:0034660) | 1.94E-03 |
| cellular component organization or biogenesis (GO:0071840) | 2.48E-03 |
| maturation of LSU-rRNA (GO:0000470) | 3.28E-03 |
| translational elongation (GO:0006414) | 4.48E-03 |
| cold acclimation (GO:0009631) | 7.73E-03 |
| regulation of translation (GO:0006417) | 9.17E-03 |

**Table 3.**
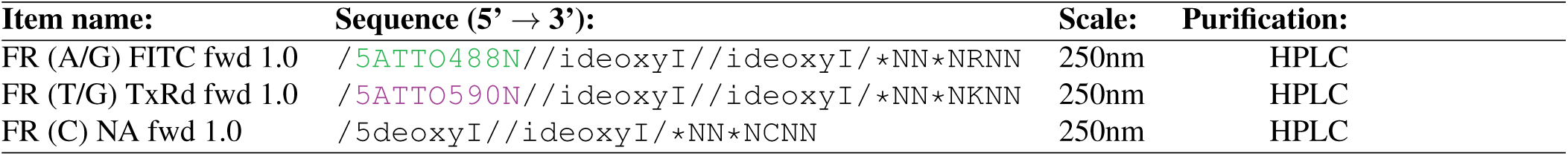
Forward Read set PRICKli probes, version 1.0.

**Table 4.** Reverse Read set PRICKli probes, version 1.0 *order as custom RNA oligos.

| Item name: | Sequence (5' → 3'): | Scale: | Purification: |
| --- | --- | --- | --- |
| RR (A/G) Cy5 rev 1.0 | /5Phos/NNrRNN/ideoxyI//ideoxyI//ideoxyI//3ATTO647NN/ | 100nmR | RNASE |
| RR (T/G) Cy3 rev 1.0 | /5Phos/NNrKNN/ideoxyI//ideoxyI//ideoxyI//3ATTO550N/ | 100nmR | RNASE |
| RR (C) NA rev 1.0 | /5Phos/NNrCNN/ideoxyI//ideoxyI//3InvdT/ | 100nmR | RNASE |

**Table 5.**
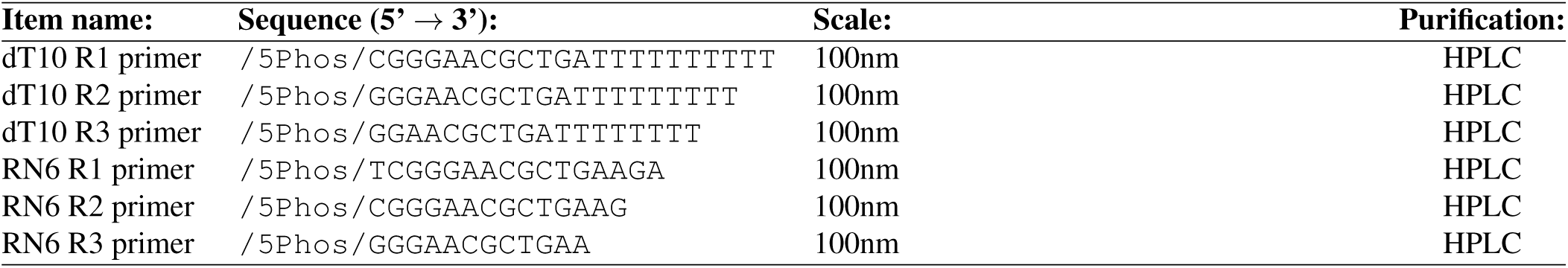
Shrinkage in situ sequencing primers, version 1.0.

**Table 6.**
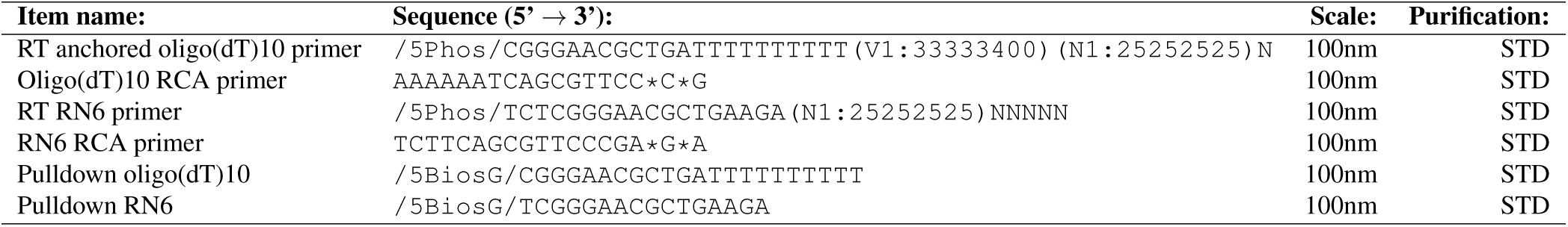
Library primers, version 1.0.

